# Macroscopic digestive anatomy of ring-tailed lemurs (*Lemur catta*), including a comparison of frozen and formalin-stored specimens

**DOI:** 10.1101/2020.07.16.206201

**Authors:** Marcus Clauss, Jelscha Trümpler, Nicole L. Ackermans, Andrew C. Kitchener, Georg Hantke, Julia Stagegaard, Tomo Takano, Yuta Shintaku, Ikki Matsuda

## Abstract

Digestive tract measurements are often considered species-specific, but little information exists on the degree to which they change during ontogeny within species. Additionally, access to anatomical material from nondomestic species is often limited, with fixed tissues possibly representing the only available source, though the degree at which this material is representative in terms of dimensions and weight is debatable. In the present study, the macroscopic digestive tract anatomy (length of intestinal portions, and tissue weights of stomach and intestines) of *n=58 Lemur catta* (from neonates to 25 years of age) was quantified, of which 27 had been stored frozen and 31 fixed in formalin. Particular attention was paid to the caecum and the possible presence of an appendix. The intraspecific allometric scaling of body mass (BM)^0.46[0.40;0.51]^ for total intestine length and BM^0.48[0.41;0.54]^ for small intestine length was higher than the expected geometric scaling of BM^0.33^, similar to literature results on interspecific scaling. This difference in scaling is usually explained by the hypothesis that the diameter of the intestinal tube cannot increase geometrically, to maintain optimal absorption. Therefore, geometric volume gain of increasing body mass is accommodated by more-than-geometric length scaling. Literature describes *L. catta* as being variable with respect to the presence of an appendix. No appendix was found in specimens of the present study. The proportions of length measurements did not change markedly during ontogeny, indicating that proportions developed in the foetus are already representative for the adult animal. By contrast, width and tissue-mass scaling of the caecum indicated a disproportionate growth of this organ during ontogeny that was not reflected in its length. Compared to overall intraspecific variation, the method of storage (frozen vs. formalin) had no relevant impact on length or weight measurements.

## Introduction

Based on geometric considerations, volume measurements should scale isometrically (in other words, linearly) with body mass, surface measurements should scale to body mass to the power of 0.67, and length measurements – such as the length of intestinal tract sections – should scale to body mass to the power of 0.33 (Calder 1996; Clauss and Hummel 2005). However, several studies found a higher scaling exponent for interspecific scaling relationships of various intestinal section lengths with body mass in mammals (Woodall and Skinner 1993; Lavin et al. 2008; McGrosky et al. 2016; McGrosky et al. 2019a; McGrosky et al. 2019b). The common explanation for this observation, developed to our knowledge by Woodall and Skinner (1993), is that on the one hand, both intestinal volume and surface area do indeed scale geometrically with body mass, but the intestinal diameter scales to a lower exponent in order to maintain short diffusion distances from the lumen to the secretive and absorptive surfaces. Therefore, the length of the intestine must scale more-than-geometrically to accommodate geometric volume and surface scaling. If this reasoning were correct, we would expect a similar scaling at the intraspecific level across ontogeny, particularly because the transition from milk to any other diet generally implies a decrease in diet digestibility, theoretically making short distances between lumen and surface all the more relevant.

On a completely different level of consideration, anatomical material from nondomestic species can be hard to come by. Given that hunting expeditions or culling operations are no longer socially acceptable, the accretion of sample sizes typically depends on storage of deceased individuals collected as single specimens from zoological collections, or during field work. Typically, this storage occurs either as frozen material, or fixed in formalin. Formalin often leads to tissue shrinkage compared to fresh material (Lentle et al. 1997), and comparisons of intestine length measurements between frozen or formalin-fixed material on a limited number of specimens indicated some degree of shortening during formalin storage (Hume et al. 1993). Most recently, a comparative study in humans in which intestine length was measured during abdominal surgery in live patients and at dissection in formalin-fixed cadavers, indicated significantly shorter length measured in the fixed specimens; the results also indicated that the length measures in the fixed specimens were shorter than those reported for freshly dissected cadavers in the literature (Zhou et al. 2020). Evidently, these results leave room for shrinkage to be a consequence of time after death itself, irrespective of the method. For example, studies on skin samples indicated that tissue shrinkage occurred as an effect of excision and was not exacerbated by formalin storage (Dauendorffer et al. 2009), and fishes shrank shortly after killing, irrespective of the preservation method, without additional effects of longer storage (Parker 1963). For intestines, both the effects of relaxation-elongation and of contraction-shortening after death have been reported literature (Zhou et al. 2020). An older comprehensive study in dogs documented intestinal shortening of fresh material occurred within the first few hours after death (Nickel 1933). This shortening sometimes persisted, but that it was more often followed by a relaxation that exceeded the shortening within 48 hours, leading to longer-than-life measurements at this timepoint that are considered to represent the relaxation of the natural tonus of the smooth intestinal musculature (Nickel 1933). These findings add to the overall uncertainty of measuring intestinal lengths, and effects of storage and fixation will depend on the state of the material at the moment of applying fixatives or freezing. There are most likely many other factors, such as whether material is frozen or fixed with or without the mesenteries, the temperature at dissection, or the forces involved in laying out an intestinal section into a straight line for measuring (Underhill 1955), that could all influence the final outcome.

In the present study, we used the opportunity of access to three different collections of gastrointestinal tracts of ring-tailed lemurs (*Lemur catta*), either preserved frozen attached to the mesenteries, or preserved in formalin after dissection of the mesenteries. The main aims of the study were to test whether intraspecific allometries of intestinal lengths resembled those reported for interspecific comparisons in other mammals, and whether a systematic difference between the two preservation methods could be detected. In addition, we aimed to investigate whether the prominence of the caecum, the site of microbial fermentation of complex carbohydrates deriving from plant fibre (Campbell et al. 2000), changed across ontogeny from milk-dependent neonates to mature individuals. Ring-tailed lemurs have been described as a species in which a caecal appendix may occur variably (Smith et al. 2013, 2017), macroscopic description of the caecum did not indicate the presence of an appendix (Campbell et al. 2000; McGrosky et al. 2019b). Therefore, special attention was given to the appearance of the apex of the caecum.

## Methods

Three different specimen collections of ring-tailed lemurs (*Lemur catta*) were available for the study, with a total of *n=58*. One consisted of 12 specimens (from neonates to 24 years old, body mass 0.07-3.48 kg, 7 females and 5 males) from various zoological collections, stored frozen as whole carcasses for varying amounts of time (3-20 years), and thawed and dissected for the present study. The second set consisted of 15 specimens (0.1-16 years of age, 0.57-2.67 kg, 6 females and 9 males) of a large family group originating from a single zoological facility, whose gastrointestinal tract had been excised immediately after death, including all mesenteries, and stored frozen until dissection (for 12 months). The third consisted of 31 specimens (known ages 0.33-25 years, 0.73-2.85 kg, 19 females and 12 males) from a single zoological collection, where the gastrointestinal tract had been dissected from freshly deceased specimens, freed of mesenteries, partially opened (along their lengths), and stored in formalin for varying amounts of time (1-60 years). In total, 18 animals surpassed the mean body mass range for free-living ring-tailed lemurs of 2-2.5 kg (Sussman 1991; Drea and Weil 2008), yet our age-body mass graph (Fig. 1) largely resembled that given by Koyama et al. (2008), and only two animals appeared to be of excessive weight for their age, being distinctively above maximum weights recorded in the natural habitat of 2.6 kg by Simmen et al. (2010). These findings suggested that obesity was not a major factor in the study populations. These two animals were excluded from all analyses. No animals of unknown age exceeded 2.60 kg.

**Figure 1.**
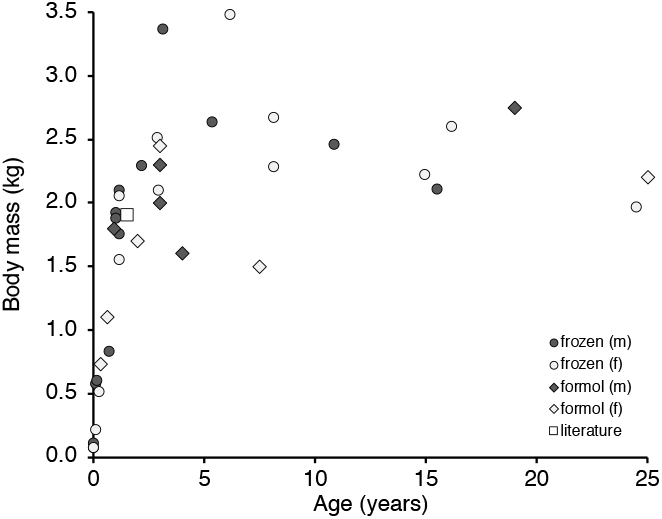
Relationship of body mass and age in the male (m) and female (f) ring-tailed lemurs (*Lemur catta*) for which age was known in the present study, separated by whether their intestine was stored frozen or in formalin. Note two particularly heavy animals at body mass > 3 kg, which were excluded from subsequent analyses.

All gastrointestinal tracts were freed from mesenteries and adhering adipose tissue and photographed (Fig. 2). For photography and measurements, thawed intestines were laid out without deliberate stretching beyond that countered by the friction of the intestine on the metal dissection table. Intestines preserved in formalin were gently pulled to a straight form for length measurements. Length measurements included the small intestine, caecum, and the combination of colon and rectum. In thawed (unopened), but not in formalin-preserved (and generally opened) caeca, the width at the base was measured as well. Subsequently, the stomach, small intestine, caecum and the joined colon-rectum were cleared of contents, blotted dry with paper towels, and weighed.

**Figure 2.**
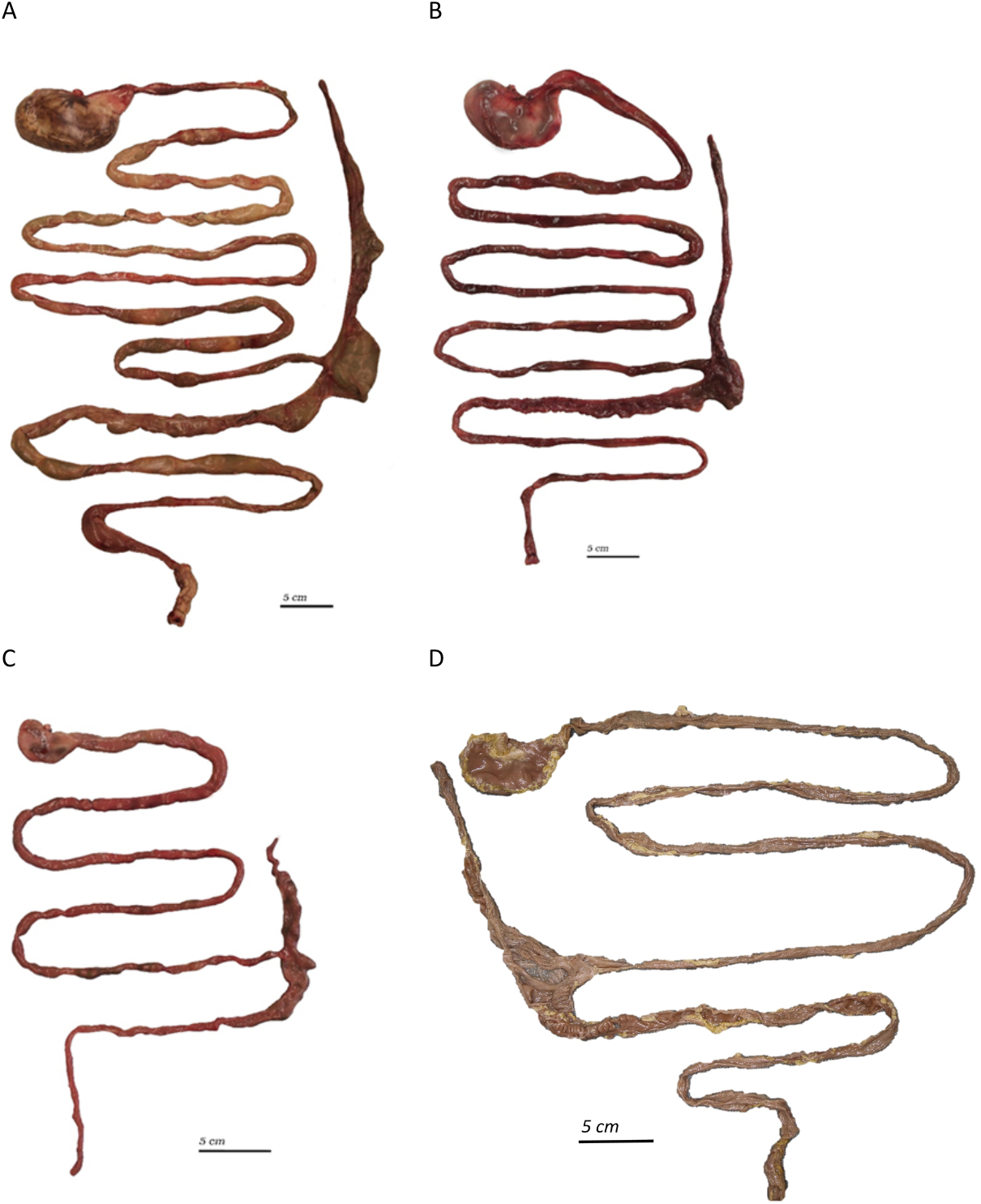
Gastrointestinal tracts of ring-tailed lemurs (*Lemur catta*) stored frozen (A-C) or in formalin (D): (A) a 16.2-year-old female of 2.6 kg, (B) an 8.2-year-old female of 2.3 kg, (C) a 0.1-year-old male of 0.6 kg, (D) an adult female of 2.1 kg.

Statistical evaluations were performed in R (R Core Team 2017). Linear models based on log-transformed data were used. First, we tested the allometric relationships of all intestine lengths, caecum width and weights with body mass. Additionally, the effect of body mass on organ measurements expressed as % of either total intestinal length or total gastrointestinal tract (GIT) tissue weight were assessed in the same manner, to test whether changes in the prominence of organs occurred across maturation. The results are given to facilitate comparison with other allometries, even though residuals of the models were mostly not normally distributed. Then, the models were repeated, including information about sex and preservation method (frozen or fixed), where applicable, including the interaction using ranked data for quantitative measures, and including body mass. Whether body mass was a significant covariable in these secondary models or not was identical to whether the scaling exponent was significant in the primary models, and is therefore not indicated separately. The significance level was set to 0.05.

## Results

The macroscopic appearance of the ring-tailed lemurs’ digestive tracts in the present study resembled that shown previously by Campbell et al. (2000) and McGrosky et al. (2019b) (Fig. 2). Although in some individuals, the visual impression of a caecal appendix appeared possible (e.g., Fig. 2C), when opened, the apex of the caecum never visually indicated a different section and did not visually resemble a lymphatic organ (Fig. 3).

**Figure 3.**
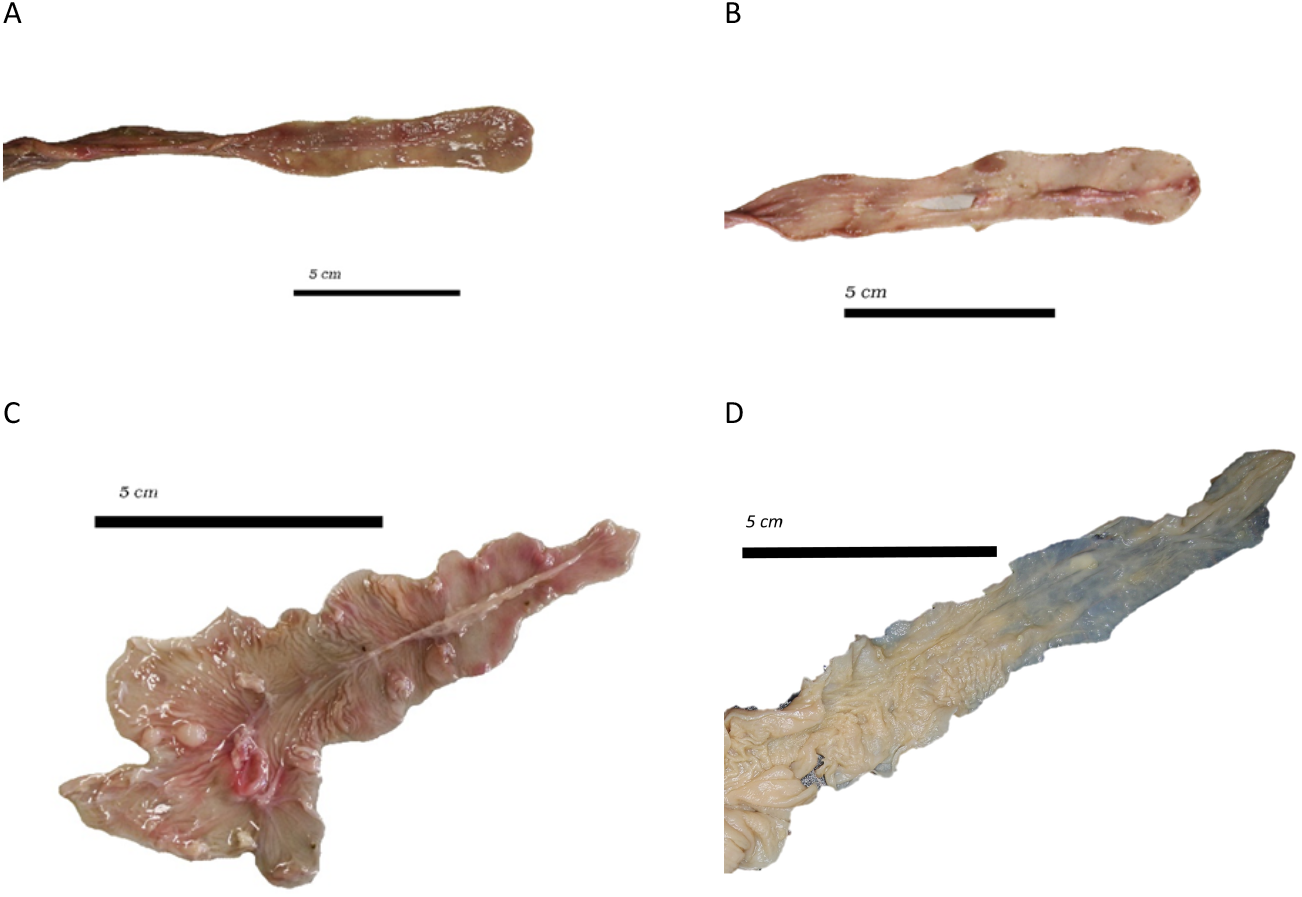
The opened caecum in various ring-tailed lemurs (*Lemur catta*), with apex pointing towards the right, suggesting an absence of a caecal appendix, stored frozen (A-C) or in formalin (D): (A) a 16.2-year-old female of 2.6 kg, (see Fig. 2A), (B) a 3.2-year-old male of 3.4 kg, (C) a 0.1-year-old male *L. catta* of 0.6 kg, (see fig. 2C), (D) an adult female of 1.5 kg.

The measurements taken in the present study were of a range that included those of an individual published by Campbell et al. (2000) (Fig. 4) except for the caecum, for which these authors reported a greater length. The allometric scaling of length measures of all intestinal sections yielded exponents whose 95% confidence intervals were above the 0.33 scaling exponent expected from geometry (Table 1). The scaling exponent of caecum width was particularly high at 0.57. The relative length of the intestinal sections did not change with body mass, suggesting that their proportions (small intestine 59%, caecum 8%, colon & rectum 32% of total intestinal length) remain stable during ontogeny (Table 1; Fig. 5A). Preservation method only had an effect on the length of the colon & rectum, where specimens preserved in formalin had higher values, but the results also indicated an interaction with sex, suggesting that preservation method affected the sexes differently in the dataset (Table 1; Fig. 4C). Therefore, samples fixed in formalin had a slightly longer relative colon & rectum length (34 ± 4 % vs. 30 ± 4 %) and a shorter relative small intestine length (57 ± 4 % vs. 62 ± 5 %) (Table 1).

**Figure 4.**
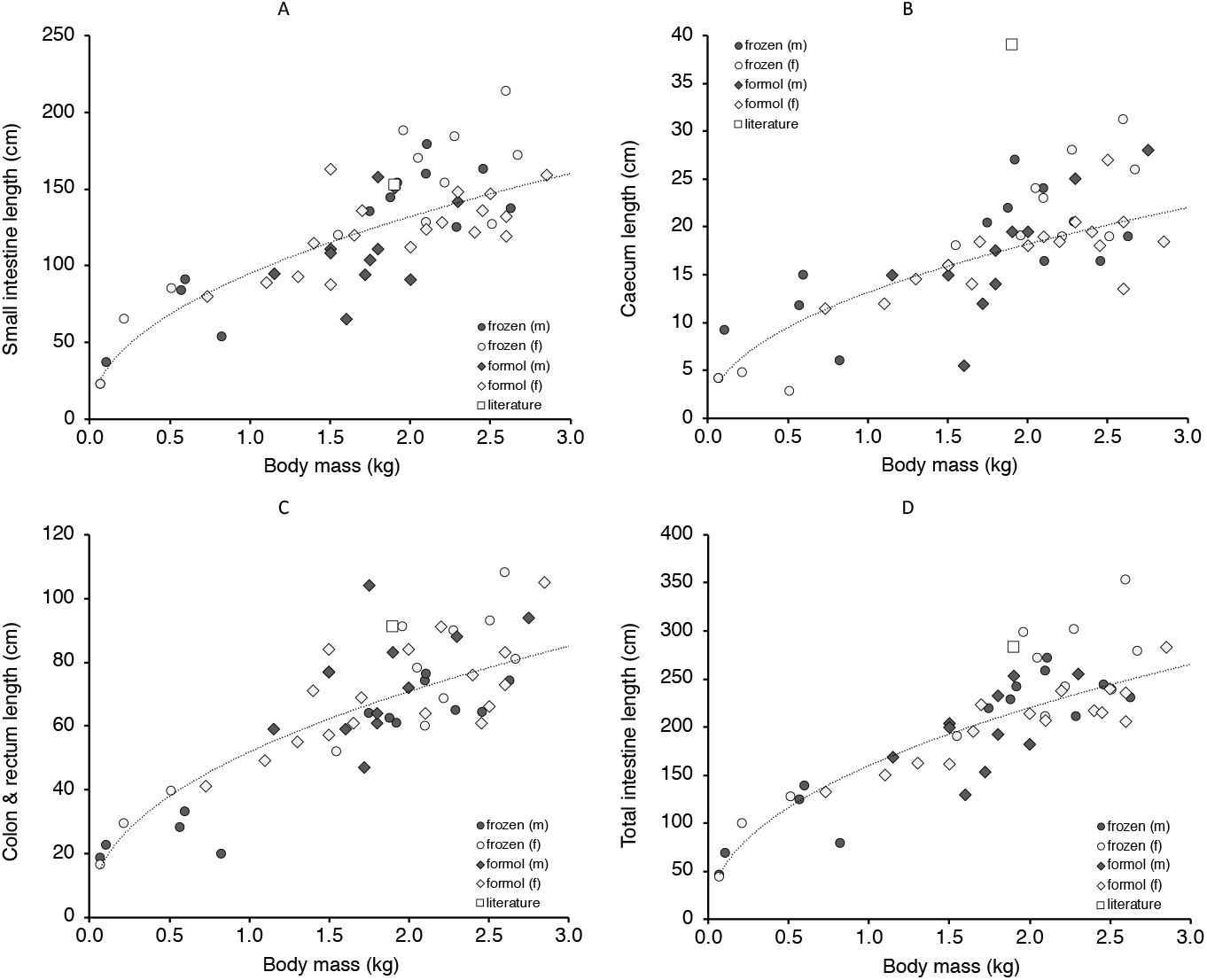
Relationship between body mass (A) small intestine, (B) caecum, (C) colon and rectum length and (D) total intestine length in ring-tailed lemurs (*Lemur catta*) of the present study (males and females, preserved frozen or in formalin), compared to an individual from Campbell et al. (2000). Statistics in Table 1.

**Figure 5.**
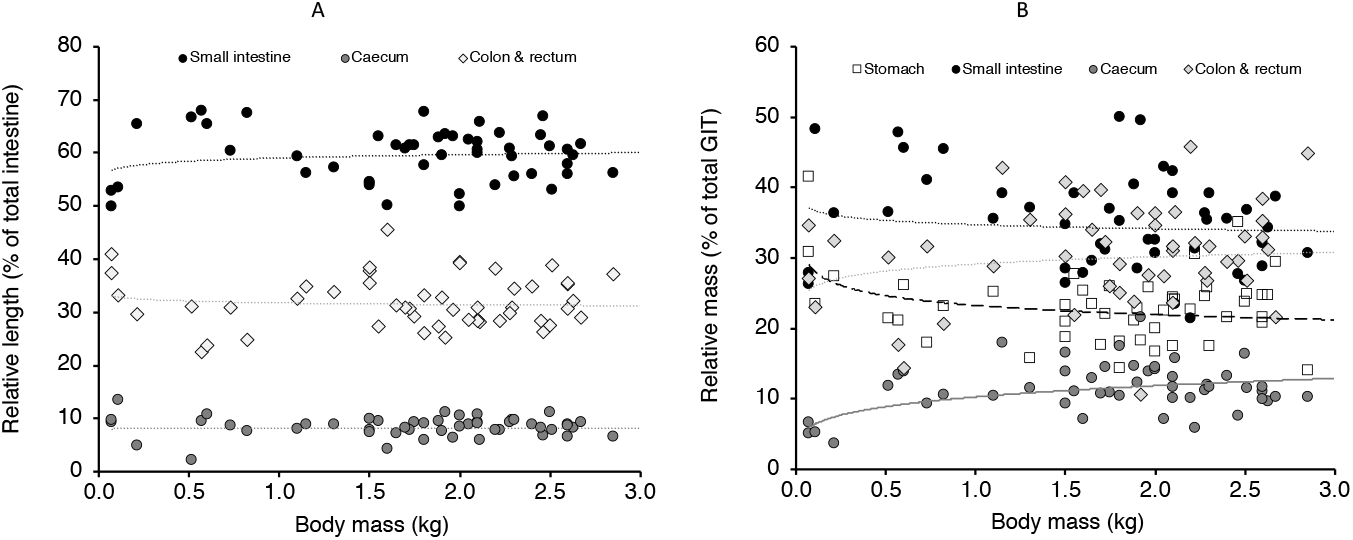
Relationship of body mass and (A) relative intestine lengths; (B) relative gastrointestinal organ masses (both in % of total) in ring-tailed lemurs (*Lemur catta*) of the present study. The only significant relationships are those for the relative mass of stomach and caecum (cf. Table 2).

**Table 1.** Allometric regressions according to y = *a* body mass^*b*^ including 95% confidence intervals for measures of intestinal lengths of ring-tailed lemurs (*Lemur catta*) of different sex and preservation method

| Dependent variable<br>(y) | *a*<br>[95% CI] | *P* | *b*<br>[95% CI] | *P* | sex<sup>1</sup> | preservation<sup>1</sup> | interaction<sup>1</sup> |
| --- | --- | --- | --- | --- | --- | --- | --- |
| *Absolute length (cm)* |  |  |  |  |  |  |  |
| Small intestine | 95 [89;101] | <0.001 | 0.48 [0.41;0.54] | <0.001 | n.s. | n.s. | n.s. |
| Caecum* | 13 [12;14] | <0.001 | 0.47 [0.37;0.58] | <0.001 | n.s. | n.s. | n.s. |
| Colon & rectum* | 52 [49;55] | <0.001 | 0.45 [0.38;0.52] | <0.001 | (+) 0.033 | (+) 0.006 | 0.028 |
| Total intestine* | 160 [152;168] | <0.001 | 0.46 [0.40;0.51] | <0.001 | n.s. | n.s. | n.s. |
| Caecum width | 2.1 [1.9;2.4] | <0.001 | 0.57 [0.47;0.67] | <0.001 | n.s. | - | - |
| *Relative length (% total intestinal length)* |  |  |  |  |  |  |  |
| Small intestine | 59 [58;61] | <0.001 | 0.01 [-0.01;0.04] | 0.282 | n.s. | n.s. | n.s. |
| Caecum* | 8 [8;9] | <0.001 | 0.01 [-0.09;0.10] | 0.913 | n.s. | n.s. | n.s. |
| Colon & rectum | 32 [30;33] | <0.001 | -0.01 [-0.06;0.03] | 0.568 | n.s. | (+) 0.014 | n.s. |

body mass in kg; \*indicates that residuals were not normally distributed
<sup>1</sup>results of additional models testing for effects of sex and preservation using ranked data; (+) indicates higher values in females, or higher values in formalin-preserved tissues

The body-mass scaling exponents for organ masses exceeded a linear scaling in their 95% confidence interval, except for the stomach (Table 2). Of the intestine sections, the small intestine showed the least distinct deviation (exponent confidence interval: 1.01;1.28), and the caecum (1.21;1.54) the most distinct deviation from linearity. As for length measures, preservation status had an effect on caecum and colon & rectum tissue mass (Table 2). Therefore, total intestinal mass and total GIT mass were slightly higher in formalin-fixed specimens (Table 2). Considering the entire GIT, the relative mass of the small intestine and colon & rectum did not vary with body mass; by contrast, the relative mass of the stomach declined, and that of the caecum increased, with body mass (Table 2; Fig. 5B). Considering only the intestinal tract, relative caecum mass increased with body mass (Table 2). Relative tissue mass of the colon and rectum of the total GIT was higher in formalin-fixed specimens (frozen: 27 ± 7 %; fixed 35 ± 6 % of total GIT mass) (Table 2).

**Table 2.**
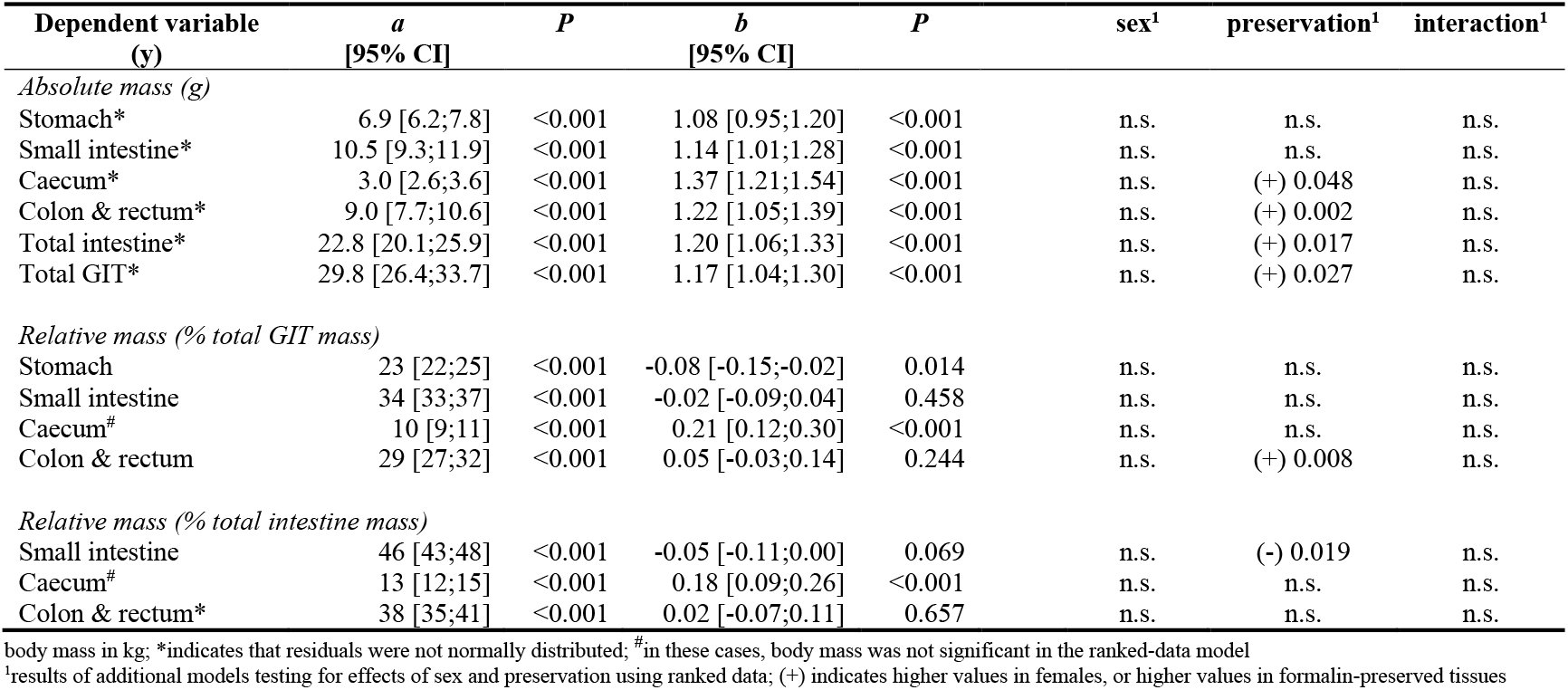
Allometric regressions for weight measurements according to y = *a* body mass^*b*^ including 95% confidence intervals for measures of tissue mass of gastrotintestinal tracts (GIT) of ring-tailed lemurs (*Lemur catta*) of different sex and preservation method

| Dependent variable<br>(y) | *a*<br>[95% CI] | *P* | *b*<br>[95% CI] | *P* | sex<sup>1</sup> | preservation<sup>1</sup> | interaction<sup>1</sup> |
| --- | --- | --- | --- | --- | --- | --- | --- |
| *Absolute mass (g)* |  |  |  |  |  |  |  |
| Stomach* | 6.9 [6.2;7.8] | <0.001 | 1.08 [0.95;1.20] | <0.001 | n.s. | n.s. | n.s. |
| Small intestine* | 10.5 [9.3;11.9] | <0.001 | 1.14 [1.01;1.28] | <0.001 | n.s. | n.s. | n.s. |
| Caecum* | 3.0 [2.6;3.6] | <0.001 | 1.37 [1.21;1.54] | <0.001 | n.s. | (+) 0.048 | n.s. |
| Colon & rectum* | 9.0 [7.7;10.6] | <0.001 | 1.22 [1.05;1.39] | <0.001 | n.s. | (+) 0.002 | n.s. |
| Total intestine* | 22.8 [20.1;25.9] | <0.001 | 1.20 [1.06;1.33] | <0.001 | n.s. | (+) 0.017 | n.s. |
| Total GIT* | 29.8 [26.4;33.7] | <0.001 | 1.17 [1.04;1.30] | <0.001 | n.s. | (+) 0.027 | n.s. |
| *Relative mass (% total GIT mass)* |  |  |  |  |  |  |  |
| Stomach | 23 [22;25] | <0.001 | -0.08 [-0.15;-0.02] | 0.014 | n.s. | n.s. | n.s. |
| Small intestine | 34 [33;37] | <0.001 | -0.02 [-0.09;0.04] | 0.458 | n.s. | n.s. | n.s. |
| Caecum<sup>#</sup> | 10 [9;11] | <0.001 | 0.21 [0.12;0.30] | <0.001 | n.s. | n.s. | n.s. |
| Colon & rectum | 29 [27;32] | <0.001 | 0.05 [-0.03;0.14] | 0.244 | n.s. | (+) 0.008 | n.s. |
| *Relative mass (% total intestine mass)* |  |  |  |  |  |  |  |
| Small intestine | 46 [43;48] | <0.001 | -0.05 [-0.11;0.00] | 0.069 | n.s. | (-) 0.019 | n.s. |
| Caecum<sup>#</sup> | 13 [12;15] | <0.001 | 0.18 [0.09;0.26] | <0.001 | n.s. | n.s. | n.s. |
| Colon & rectum* | 38 [35;41] | <0.001 | 0.02 [-0.07;0.11] | 0.657 | n.s. | n.s. | n.s. |

body mass in kg; \*indicates that residuals were not normally distributed; <sup>#</sup>in these cases, body mass was not significant in the ranked-data model
<sup>1</sup>results of additional models testing for effects of sex and preservation using ranked data; (+) indicates higher values in females, or higher values in formalin-preserved tissues

## Discussion

The present study has relevance for both methodological and biological aspects of digestive tract anatomy. The data indicate substantial intraspecific variation in intestinal measurements within mature specimens. For example, Fig. 4D indicates that at 2 kg of body mass, the length of the intestinal tract in ring-tailed lemurs may vary by a metre. In humans, at evidently higher body masses, the documented variation in small intestine length can be more than four metres (reviewed in Zhou et al. 2020). A large body of literature exists that documents intraspecific variation in intestinal length in rodents due to diet and/or energetic constraints, and intraspecific intestinal length flexibility has been linked to the number of different habitats small rodent species can occupy (Naya et al. 2008). However, for large mammals no corresponding compilations exist. To date, variation such as described in humans or in the lemurs of the present study remains largely unexplained. In the lemurs, effects of diet or different husbandry conditions appear unlikely. Thus, the variation remains unexplained. While such variation may not be a systematic problem for large-scale comparisons of intestinal length (Woodall and Skinner 1993; Lavin et al. 2008; Lovegrove 2010), studies exploring quantitative differences between only a few specimens of a few species need to take this variation into account and should include a sufficient number of individuals.

Given the magnitude of this general intraspecific variation, the variation introduced by the use of formalin-fixed material appeared to be of a negligible magnitude in the present study. The material that was compared was either frozen at unknown times (but most likely within 24 hours) after death, then thawed and dissected from the mesenteries, or dissected at unknown times after death and subsequently placed in formalin. Of the two processes, freezing and thawing could be assumed to counteract any potential effect of post-mortem contraction. By contrast, formalin fixation could theoretically have occurred at any stage of post-mortem contraction or relaxation, and therefore, on average shorter dimensions could have been expected for this method. However, if at all, the opposite was the case, with formalin-fixed specimens showing somewhat longer large intestines (Table 1). One theoretical explanation could be the effect of opening the intestines lengthwise, with no effect on the length of the smooth-walled small intestine, but with an effect on the haustrated large intestine: if opened, the haustra might not constrain the length of the organ as much as in a closed state. Unfortunately, this finding only became evident after the frozen/thawed material had been disposed, otherwise a comparison of the length of the same material with a closed and an opened intestine could have been performed. However, the formalin-fixed large intestines were also heavier – and longitudinal cuts should not affect mass measurements – suggesting that this difference between the preservation methods for the large intestine might simply have been due to chance.

Regardless of the large variation in intestinal measures in mature specimens, the intraspecific allometry, including neonates and juveniles, yielded scaling relationships comparable to those previously reported in the literature in interspecific studies (see Introduction), in the range of a 0.4-0.5 scaling exponent. A similar, more-than-geometric intraspecific scaling of the small and the large intestine across ontogeny was demonstrated in rats (Toloza and Diamond 1992) and mice (Wołczuk et al. 2011). As in the ring-tailed lemurs, the scaling effect was mostly found in the neonate and juvenile stages, and was not evident within the mature specimens. The more-than-geometric scaling of intestinal lengths, as explained in the introduction, appears to be a general feature of mammalian macroanatomy.

In ruminants and possibly other foregut fermenters, the change in proportions of the different GIT sections when going from milk-feeding to weaning, are very distinct. Indeed the fermentation compartments increase disproportionately in tissue weight (e.g., Wardrop and Coombe 1960; Godfrey 1961) and, by inference, in volume. In horses, the length proportion of the caecum and proximal colon – which represent the fermentation chambers – similarly increase with age until maturity (Smyth 1988). For the ring-tailed lemurs, a similar ontogenetic change in GIT proportions linked to the change in diet was not evident in length measurements.

The scaling of organ tissue masses surprisingly appeared more-than-linear, with 95% confidence intervals of the scaling exponent consistently above 1.00 (Table 2). To our knowledge, no recent comprehensive interspecific treatise for gastrointestinal tissue mass exists. Calder (1996) cites the scaling in 41 mammal species established by Brody (1945) with an exponent of 0.94; using the standard error for the exponent given in the original by Brody (1945), the 95% confidence interval of that exponent includes linearity at 0.85-1.03 (and is nearly identical for the exponent found for birds in that study). In the original data from Navarrete et al. (2011) for 100 mammal species, a similar scaling exponent with a confidence interval of 0.88-0.94 can be calculated, and Prothero (2015) found a scaling exponent of 0.93 in mammals that also excluded linearity in the confidence interval. Why mammalian and avian GIT scaling should be slightly less-than-linear has not been explained so far, and we also do not offer an explanation. We hypothesize that the more-than-linear scaling found in our data is an intraspecific effect of ontogeny, reflecting the shift from milk feeding to solid food. In the case of the ring-tailed lemur, the natural diet comprises fruits, leaves and other plant parts (e.g., Rasamimanana and Rafidinarivo 1993; Simmen et al. 2006). On the one hand, an increasing tissue mass with age could derive from a disproportionately increased muscle mass as an effect of processing solid material. On the other hand, it could derive in particular from absorptive mucosa development in those compartments (caecum, colon) where fermentative digestion intensifies after the switch to solid food. The enhancing effect of short-chain fatty acids, the main products of microbial fermentation, on gut mucosa development - and hence tissue mass - is well-known (e.g., Kripke et al. 1989). Fermentative microbial digestion has been suggested for ring-tailed lemurs (Campbell et al. 2000) and was demonstrated by the measurement of short-chain fatty acids in the faeces of captive specimens (McKenney et al. 2018). Correspondingly, the scaling exponent of tissue mass was highest for the caecum, followed by the colon and rectum, whereas the small intestine only scaled slightly higher than linearly, and the stomach scaling was linear (Table 2). A shift in the faecal microbiome from milk-feeding to weaning has been demonstrated in lemurs, including ring-tailed lemurs (McKenney et al. 2018), which would be expected to parallel the increased tissue mass.

The present study did not find evidence for the presence of a caecal appendix in ring-tailed lemurs. The review of the primate appendix by Fisher (2000) did not include ring-tailed lemurs as either a species with or without an appendix, and neither Campbell et al. (2000) nor McGrosky et al. (2019b) reported evidence for an appendix in ring-tailed lemurs. The external appearance of the caeca of some individuals included an apparent narrowing of the caecal apex that created the impression of an appendix (Fig. 2), but neither thickening of the mucosa nor macroscopic appearance of lymphatic tissue were evident (Fig. 3). A recent description of the gastrointestinal anatomy of another lemur species, *Eulemur coronatus*, also did not suggest the presence of an appendix (Schwitzer 2009), although the species is among those for which an appendix is assumed in the literature (Fisher 2000; Smith et al. 2013, 2017). A more detailed histological study of the putative appendices of lemur species might be interesting.

## Acknowledgements

We thank the contributing zoological collections for donating the material for scientific curation. We thank the Japan Monkey Centre for facilitating the project and sharing their knowledge of the lemurs. This study (Japanese samples) was conducted in compliance with guidelines for care and use of nonhuman primates by the Japan Monkey Centre. This study was partly financed by JSPS KAKENHI (#19H03308 to IM). National Museums Scotland thanks the Negaunee Foundation for their generous support of a curatorial preparator.

